# Motor output and control input in flapping flight: a compact model of the deforming wing kinematics of manoeuvring hoverflies

**DOI:** 10.1101/759175

**Authors:** Indira Nagesh, Simon M. Walker, Graham K. Taylor

## Abstract

Insects are conventionally modelled as controlling flight by varying a few summary kinematic parameters that are defined on a per-wingbeat basis, such as the stroke amplitude, mean stroke angle, and mean wing pitch angle. Nevertheless, as insects have tens of flight muscles and vary their kinematics continuously, the true dimension of their control input subspace is likely to be much higher. Here we present a compact description of the deforming wing kinematics of 36 manoeuvring *Eristalis* hoverflies, applying functional principal components analysis to Fourier series fits of the wingtip position and wing twist measured over 26,541 wingbeats. This analysis offers a high degree of data reduction, in addition to insight into the natural kinematic couplings. We used statistical resampling techniques to verify that the principal components were repeatable features of the data, and analysed their coefficient vectors to provide insight into the form of these natural couplings. Conceptually, the dominant principal components provide a natural set of control input variables that span the control input subspace of this species, but they can also be thought of as output states of the flight motor. This functional description of the wing kinematics is appropriate to modelling insect flight as a form of limit cycle control.

## 1. Introduction

Insects in general, and flies in particular, display a spectacular degree of manoeuvrability, which they achieve despite—or perhaps because of—their lack of independent control surfaces analogous to the flaps of a conventional fixed-wing aircraft or the tail of a bird (but see [1]). Instead, the kinematics of their beating wings are continuously adjusted to meet the joint requirements of weight support, thrust production, and flight control simultaneously. In this respect, insects resemble a helicopter with an articulated rotor, but their wing kinematics are infinitely more complex and varied than those of a rotor blade, even though both function as passive aeroelastic structures driven and controlled by torques applied at the wing root. This difference ultimately reflects the complexity of the underlying drive mechanism. In particular, the torques that are applied at the root of an insect’s wing are produced by a multitude of different flight muscles contained within the thorax [2].

The various flight muscles may be broadly categorised into two types: power muscles, and steering muscles. In most insects, including dipteran flies, the main power muscles do not attach directly to the wings, but instead drive the wingbeat by deforming the thorax, through which mechanical energy is propagated to the wing hinge [3]. In Diptera, and several other insect orders, these indirect muscles are asynchronous and stress-activated, which means that their neuronal activity does not map onto their contractile state in a one-on-one fashion. Instead, as the indirect flight muscles always occur in antagonistic sets, the contraction of one set stretches the other, thereby stimulating its contraction and giving rise to a limit cycle oscillation [4]. This arrangement presents limited opportunities for the direct control of power muscle output, but because the thorax is a complex and highly anisotropic three-dimensional structure, even the simple linear motions of the indirect flight muscles are transformed into a complex three-dimensional motion after transmission through the wing hinge [3].

In contrast to the power muscles, all of the steering muscles are activated neuronally. Most of the steering muscles attach directly to the wing hinge, or close to it, controlling its complex three-dimensional motion through a variety of highly nonlinear mechanisms [2,5]. Other indirect steering muscles act by modifying the deformation of the thorax. Besides the obvious function of the direct steering muscles in changing the trajectory of the wing tip, or altering the amplitude and phase of the wing’s pitching rotation and torsional deformation, indirect steering muscle activity may be important in controlling the frequency of the limit cycle oscillations of the stretch-activated power muscles [4]. In summary, the power muscles can be thought of as driving a limit cycle oscillation of the wing; the form and frequency of which is controlled by the steering muscles [6],[4]. This perspective is consistent with the scheme proposed by [7], according to which the coupled motion of the insect’s wings and body is treated as a limit cycle oscillation that must be controlled to stabilize and steer its flight.

Given that the wings are passive aeroelastic structures, the complexity of their kinematics ultimately derives from the complexity of their actuation and deformation, with the latter in turn being shaped by the complexity of the wings’ venation [8–12]. The number and arrangement of the flight muscles varies across insects, but dipteran flies possess some 13 pairs of direct steering muscles in addition to the many other pairs of indirect power muscles and indirect steering muscles that together force the deformation of the thorax. The result is a highly coupled, nonlinear, and over-actuated system [2]. It is an open question whether the direct steering muscles can be controlled fully independently, but descending neurons from the brain are connected to more than one flight muscle in flies. Moreover, in hawkmoths, control signals from the central nervous system are distributed to multiple subunits of the indirect power muscles, leading to a synergistic action of the muscles of the left and right wings [13].

Any complete quantitative description of an insect’s wing kinematics must necessarily reflect the complexity of this underlying mechanism. Nevertheless, most previous studies have simply used a set of summary kinematic parameters defined on a per-wingbeat basis, such as the stroke amplitude or inclination of the stroke plane [14]. Evidently, these few parameters are not sufficient to capture either the dynamics of the underlying mechanism, or the detailed time history of the insect’s wingbeat. This is problematic, because the dimension of an insect’s control input subspace may therefore be far greater than has been captured previously. For instance, previous work assessing the controllability of insect flight has used 3 or 4 summary kinematic parameters defining the stroke amplitude, mean stroke angle, and mean pitch angle of the wing on the upstroke and downstroke [15,16]. This simplified approach is reasonable where the aim is simply to test whether these kinematic parameters are sufficient to stabilize flapping flight [15,16], but will not characterise fully the controllability of the real insect. Moreover, as these summary kinematic parameters are defined on a per-wingbeat basis, they cannot capture the fine details of the wing’s motion, such as whether the wing tip traces a U-shaped or figure-of-eight trajectory [17], which can have large effects on aerodynamic force production [18]. In summary, previous studies of insect flight dynamics and control have considered kinematic parameters that are easy to quantify, rather than kinematic parameters that reflect what the insect actually controls. This paper aims to rectify this position by providing a natural method of modelling an insect’s deforming wing kinematics.

Previous research has used a spherical coordinate system [17] to describe the time history of the wing tip kinematics, and has used blade-element modelling to describe the twisting of the wings through the stroke [19]. In principle, a set of 4 time-varying primary kinematic variables, comprising 2 spherical angular coordinates, and 2 coefficients measuring the spanwise twist gradient and twist offset, is sufficient to describe the motions of a flapping wing with time-varying linear twist distribution [1]. In hoverflies, the angular deflection of the alula, which is a small hinged flap at the base of the wing, represents an additional state of the system. We do not include the state of the alula in our modelling, however, because this has only previously been recorded as a binary variable [1], has a comparatively small aerodynamic effect [20], and is already known to be closely correlated with the primary wing kinematics [1]. Previous work has used these same spherical and blade-element coordinate systems as a basis for defining the wing kinematic inputs that are important in flight control. Hence, insects are usually said to control their flight by varying their stroke amplitude (i.e. the peak-to-peak amplitude of the combined variation in stroke angle and deviation angle), or their stroke plane angle (i.e. the slope of the path traced by the wingtip), or the timing of their wing rotation (i.e. the phase of the pitching motion of the wing with respect to its stroking motion) [14]. However, whilst insects must certainly be capable of exerting control over all of these aspects of their kinematics, it is an open question whether they are capable of controlling them independently, or whether the same variables come close to providing a natural representation of the insect’s control inputs.

The approach that we apply here is quite different to previous studies of insect flight, but similar in some respects to a groundbreaking study of bat wing kinematics that used principal components analysis (PCA) to quantify the dimensional complexity of the wing’s motion [21]. The wing of a bat is very different to the wing of an insect, comprising a directly actuated skeletal structure with an elastic membrane that is held under muscular tension. Nevertheless, 95% of the complexity of a bat’s wing motion could be captured by the first 16 principal components (PCs) of the three-dimensional positions of the 17 tracked markers through time, or by the first 15 PCs of the 20 corresponding joint angles [21]. This is broadly similar to the approach that we apply here, with the key difference that we undertake the PCA not on marker positions or joint angles, but rather on the Fourier series coefficients of the four primary kinematic variables that we use to describe the time history of each wingbeat. This approach to data reduction is known as functional principal components analysis (FPCA), because the PCs that the analysis identifies are expressed in terms of a set of basis functions, here comprising the terms of the truncated Fourier series. This provides an exceedingly compact description of the insect’s wing kinematics, and offers a clear way of expressing the control couplings that are observed during free-flight manoeuvres.

The analysis that we present here is based on a published empirical dataset describing the deforming wing kinematics of free-flying drone flies over a total of *N* = 26, 541 wingbeat pairs [1]. We begin by giving a brief description of the videogrammetric method already used to measure the complete time history of the 4 primary kinematic variables through each wingbeat. We then describe the analytical methods that we develop here, starting with a description of how we used periodic regression to decompose the time history of all 4 primary kinematic variables into a time-periodic component and a time-linear component for each wingbeat. We next describe how we applied FPCA to the time-periodic and time-linear components of the Fourier series that we fitted to each wingbeat, identifying the minimum number of PCs needed to explain at least 99% of the between-wingbeat variation. To the extent that this reduced set of PCs encompasses all of the observed kinematic scope of the species, it necessarily spans all of the insect’s control input subspace, thereby providing a small set of variables capturing all of the relevant variation in wing kinematics. Finally, we analyse the dominant PCs so as to characterise the key couplings that are present in the wing kinematics, before discussing how the PC decomposition provides a natural basis for defining the control input vector in a state-space model of the insect’s flight dynamics and control.

## 2. Materials and methods

The empirical measurements that we analyse here are those previously described in [1]. For completeness, we begin by providing a brief summary of the experimental methods and kinematic reconstruction technique, before providing a detailed description of the analysis that forms the primary contribution of this paper.

### (a) Experimental methods and kinematic reconstruction

A total of 36 wild-caught *Eristalis tenax* or *E. pertinax* were allowed to fly freely inside a 1m diameter opaque acrylic sphere with diffuse overhead lighting. High-speed video sequences were captured at 3.8kHz using four synchronized Photron SA3 cameras (Photron Ltd, West Wycombe, UK) with 180mm macro lenses (Sigma Imaging Ltd, Welwyn Garden City, UK) viewing the insect through clear portholes in the upper hemisphere. The recordings were triggered automatically by an infrared trigger system as the insect passed through the centre of the sphere. Two 200W pulsed infrared lasers (HSI-5000, Oxford Lasers Ltd, Oxford, UK) provided intense 805nm back-illumination invisible to the insect, with a 20*μ*s pulse duration slaved to the cameras. The cameras were calibrated using a nonlinear least squares bundle adjustment routine implemented in MATLAB, given images of a calibration grid in a wide range of positions and orientations [22]. A total of 854 video sequences were obtained in which both wings were visible for two or more wingbeats, comprising *N* = 26, 541 wingbeat pairs [1]. A typical flight sequence consisted of slow forward flight punctuated by the fast saccadic manoeuvres typical of hoverflies [23,24]. The recordings also include backward, hovering, and fast forward flight [1]. The maximum speed that we recorded was around 1 ms^−1^, with maximum horizontal accelerations approaching 1*g*.

A voxel-carving method was used to delimit the wing and body in three dimensions (see [1] for details). The primary wing kinematic parameters were defined with respect to a right-handed body-fixed axis system for the right wing, and with respect to an equivalent left-handed axis system for the left wing. Each axis system had its origin at the wing base, with the *x*-axis directed anteriorly parallel to the major axis of the body voxels, and the *y*-axis directed toward the wing tip along the line connecting the wing bases. This convention ensures equivalency by making the sign of all the primary wing kinematic parameters the same for both wings in symmetric flight. The wing tip kinematics are described by a pair of spherical coordinates measuring the azimuth (stroke angle, *ϕ*) and elevation (deviation angle, *θ*) of the wingtip. Note that because the longitudinal body axis is at an oblique angle to the stroke plane, these quantities vary to a similar degree through the wingbeat, which is different to other conventions in which the stroke angle and deviation angle are defined with respect to the stroke plane. The local pitch angle *ω*(*r*) of each wing is modelled as a linear function of the relative distance from the wing base (*r*) expressed as a proportion of the wing length, such that *ω*(*r*) = *ω*_0_ + *ω*_*r*_*r*, where the offset term is called the twist constant (*ω*_0_), and the slope term is called the twist gradient (*ω*_*r*_).

Because wingbeat frequency varied considerably between individuals (mean ± SD: 188 ± 14 Hz for all wingbeats), there was a similarly variable number of video frames recorded per wingbeat. Hence, to provide consistency in our representation of the kinematic data, we first fitted a quintic smoothing spline to our measurements of each of the four kinematic variables *ϕ*(*t*), *θ*(*t*), *ω*_0_(*t*), and *ω*_*r*_(*t*), where *t* is time. We used MATLAB to fit these splines at a tolerance calculated to remove the same total amount of variation as a 3rd order Butterworth filter with a −3dB cutoff frequency of 500Hz for the wing tip kinematics *ϕ*(*t*) and *θ*(*t*), and a −3dB cutoff frequency of 800Hz for the wing twist kinematics *ω*_0_(*t*) and *ω*_*r*_(*t*). Finally, we transformed time *t* to wingbeat phase *φ*, and evaluated the splines analytically at 101 evenly spaced points through the wingbeat on the interval *φ* ∈ [0, 2*π*], keeping separate record of the wingbeat period. We defined the phase of the wingbeat (*φ*) relative to the timing of the reversal of the mean angular velocity of the left and right wingtips at the top of the downstroke, where *φ* = 0 corresponds to the start of the downstroke. The justification for defining the phase of the wingbeat with respect to stroke reversal rather than with respect to wing twist is that the acceleration of the wing tip depends closely on the torques applied at the base of the wing, whereas wing torsion and rotation are dominated by aeroelastic forces.

We call the resulting estimates of the 4 primary kinematic variables (i.e. *ϕ*, *θ*, *ω*_0_, and *ω*_*r*_) the full-fidelity kinematics, to distinguish these from the high-fidelity analytical approximations that we arrive at through the progressive rounds of model reduction described in the next two subsections. The estimates of the four primary kinematic variables are provided as Supplementary Data 1, together with the code implementing the Fourier series analysis and the FPCA to produce the high-fidelity approximations.

### (b) Fourier series representation of the wingbeat kinematics

The first step in our reduction of these data was to express the full-fidelity kinematics, comprising 4 × (101 − 1) = 400 distinct sample points per wingbeat, as periodic functions of the wingbeat phase *φ*. We did this by using periodic regression to fit a truncated Fourier series to each of the four kinematic variables, using a 5th order Fourier series to capture the variation in the wing tip kinematics *ϕ*(*φ*) and *θ*(*φ*), and a 3rd order Fourier series to capture the variation in the wing twist kinematics *ω*_0_(*φ*) and *ω*_*r*_(*φ*). We also fitted a time-linear component for each wingbeat to account for the fact that no actual wingbeat cycle is ever strictly periodic, as in practice the wing is never found in exactly the same kinematic state at *φ* = 0 and *φ* = 2*π*. The order of the Fourier series was matched to the degree of smoothing of the splines fitted in the preceding section, and captured > 99.9% of the variation in the full fidelity kinematics through each wingbeat (mean *R*^2^ > 0.999, over all *N* = 26, 541 wingbeat pairs). The data and code required to perform this decomposition are provided as Supplementary Data 1, and a graphical portrayal of the scope of the modelling is shown for one representative wingbeat in Figure 1A-D.

**Figure 1.**
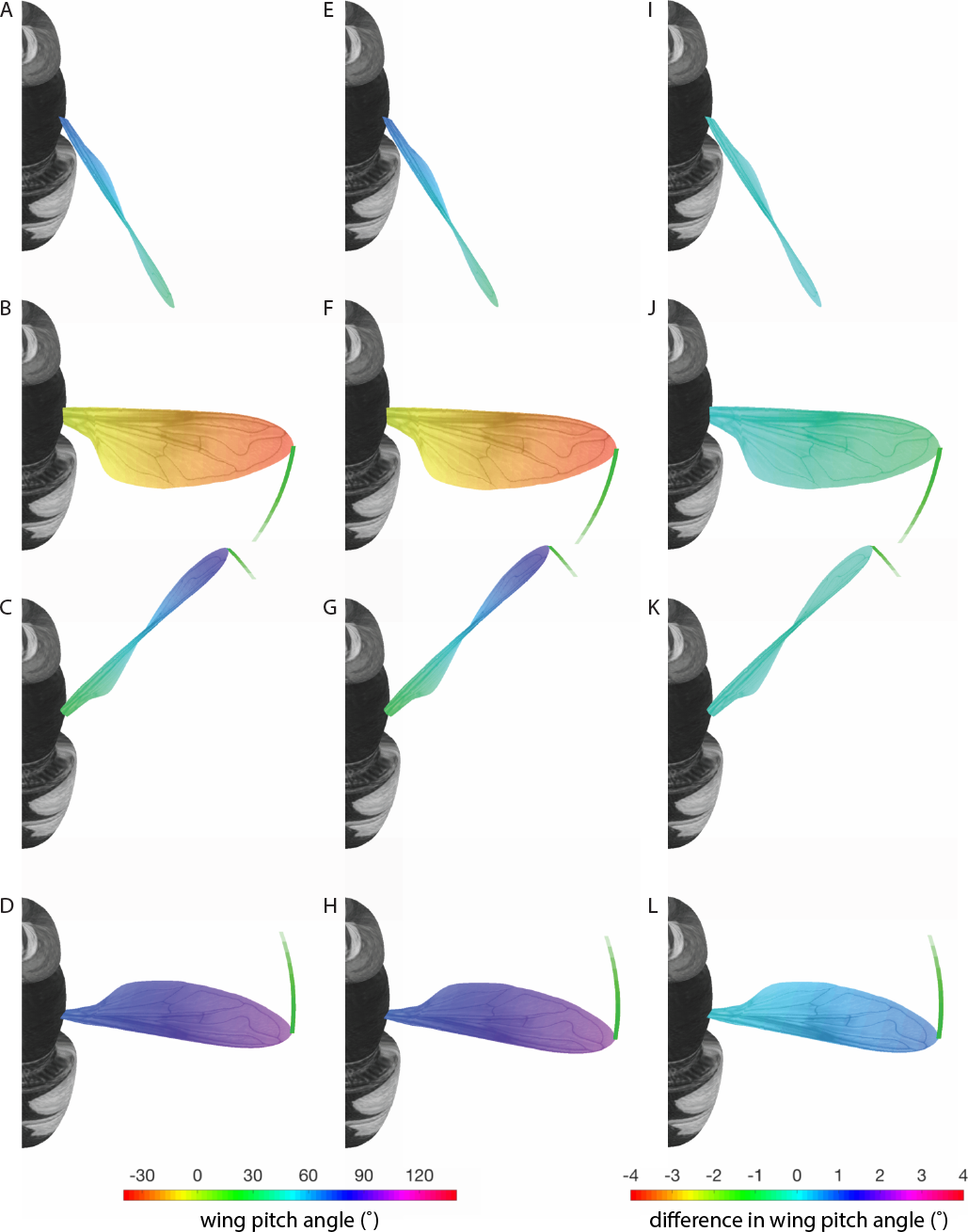
Kinematic reconstruction of a randomly selected wingbeat. (A-D) Kinematics reconstructed using all of the fitted Fourier series terms. (E-H) Kinematics reconstructed using only the first 20 time-periodic PCs and all 4 time-linear PCs. (I-L) Difference in local pitch wing angle of wing between the reduced-order PC model in (E-H) and the full Fourier series model in (I-L). Each column shows four different phases of the stroke: (A,E,I) start of downstroke, *φ* = 0; (B,F,J) mid-downstroke, *φ* = *π*/2; (C,G,K) start of downstroke, *φ* = *π*; (D,H,L) mid-upstroke, *φ* = 3*π*/2. Note the high-fidelity of the approximation at all phases of the wingbeat. Green traces show the prior wing tip trajectory. See Supplementary Video 1 for an animated version of the second and third columns of this figure, showing the effect of varying the number of time-periodic PCs used in the reconstruction.

The generic Fourier series representation for each primary kinematic parameter is:

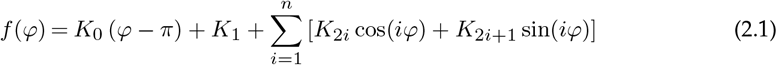

where *f* (*φ*) stands for any of the 4 kinematic variables (i.e. *ϕ*, *θ*, *ω*_0_, or *ω*_*r*_) as a function of the wingbeat phase *φ*. Breaking this expression down into its component parts, the coefficient *K*_0_ models the time-linear variation in *f* (*φ*) through each wingbeat, and is multiplied by (*φ − π*) rather than by *φ*, to ensure that this time-linear term has zero mean on the interval *φ* ∈ [0, 2*π*]. The constant offset term *K*_1_ represents the mean of *f* (*φ*) over the wingbeat, and the remaining coefficients *K*_2_ … *K*_2*n*+1_ capture the periodic variation in *f* (*φ*) through the wingbeat, where *n* is the order of the harmonics, which takes the value *n* = 3 for the wing tip kinematics and *n* = 5 for the wing twist kinematics. Analytically, these Fourier series coefficients *K*_2_ … *K*_2*n*+1_ describe a periodic oscillation about the reference condition described by the constant offset term *K*_1_. Conceptually, these time-periodic components may be thought of as representing a limit cycle oscillation of a nonlinear system, in contrast to the aperiodic motion described by the time-linear term *K*_0_, which represents the motion required to transition smoothly between different limit cycles [4,6,7].

Each wingbeat is therefore described by a total of 8 + 8 + 12 + 12 = 40 numerical coefficients. Collecting the periodic coefficients for a given wingbeat together as:

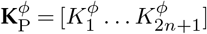

where 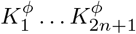 denotes the set of coefficients for the kinematic variable *ϕ*, and similarly for the other 3 kinematic variables, we may compactly summarise the entire dataset in the matrices:

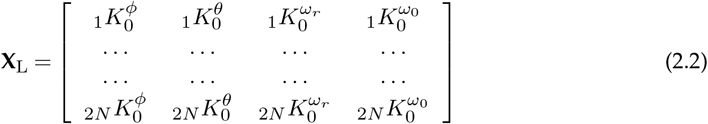

and

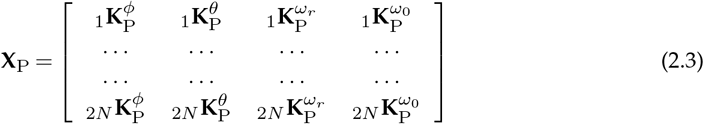

where **X**_P_ ∈ ℝ^2*N* × 36^ describes the time-periodic components, and **X**_L_ ∈ ℝ^2*N* × 4^ the time-linear components, for all *N* = 26, 541 wingbeat pairs. For a complete dimensional description of a given wingbeat, it is of course necessary to specify its period, but we did not model the variation in wingbeat frequency explicitly in our analysis. Including the wingbeat period as a variable in the subsequent FPCA would have been problematic, because its dimensions are different to those of the kinematic coefficients, all of which are in units of angular measure. This is no great limitation in practice, because whereas wingbeat period varies substantially between individuals, the within-individual variation in wingbeat period during any given flight recording was small (mean coefficient of variation: 2.3%). Because the primary kinematic variables are all in units of angular measure, and because their range of variation was similar (approximately ±50° in each case), we did not normalise any of the time-periodic or time-linear coefficients before running the FPCA.

In the next subsection, we use FPCA to describe separately the kinematic couplings that are present in the data matrices **X**_L_ and **X**_P_. This separation is appropriate because the time-linear components can be conceptualised as transition terms that allow the insect to transition from one steady state oscillation to another, and the modelling ought therefore to make allowance for these transitions to be reversible. For example, a time-periodic oscillation A should be able to transition to a time-periodic oscillation B through the superimposition of a time-linear transition C on A; conversely, this time-periodic oscillation B should be able to transition back to the same time-periodic oscillation A by superimposing a reversal of the time-linear transition C on A. This being so, there is no particular reason to expect any straightforward correlation between the time-periodic and time-linear components of a wingbeat. A more general way of conceptualising this separation is in terms of limit cycle control: the principal components of the time-periodic coefficients capture how the steady-state oscillations of the wing vary (i.e. they capture variation in the shape of the orbit in state space); the principal components of the time-linear coefficients capture how the wing transitions between different steady-state oscillations (i.e. they capture variation in the position of the orbit in state space).

### (c) Functional Principal Component Analysis (FPCA)

PCA serves to decompose a data matrix **X** ∈ ℝ^*p*×*m*^ containing *p* observations of *m* variables into the product of two new matrices:

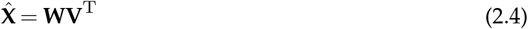

where the hat notation is used to indicate that the data matrix **X** has been centered by subtracting the mean of each column, and where the superscript T denotes the matrix transpose. The left matrix **W** is a *p* × *m* matrix of PC scores, with a typically large number of rows corresponding to the individual observations, and a typically smaller number of columns corresponding to the individual PCs. The right matrix **V**^T^ is an *m* × *m* matrix of PC coefficients, ordered such that the first column of **V** contains the PC explaining the most variation in **X**, the second the next most variation in **X**, and so on. In the special case that the observations contained in the data matrix **X** are the coefficients of a set of basis functions, as is the case for the matrices **X**_P_ and **X**_L_ (see Eqs. 2.2–2.3), then the resulting matrix decomposition is called a functional principal components analysis (FPCA).

In practice, the decomposition of a centered matrix 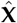 into its corresponding parts **W** and **V**^T^ is usually accomplished using the well-known singular value decomposition (SVD):

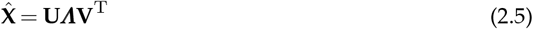

where ***Λ*** = [λ_1_, λ_2_, …, λ_*m*_] is a diagonal matrix containing the ordered eigenvalues of **X**, where **V** is the right eigenvector matrix of **X**, and where **U** is the corresponding left eigenvector matrix. By an obvious use of notation, it follows that the 2*N* × 4 matrix **W**_L_ = **U**_L_***Λ***_L_ and the 2*N* × 36 matrix **W**_P_ = **U**_P_***Λ***_P_ contain the scores for the left and right wings of all *N* = 26, 541 wingbeat pairs on the PCs identified for the centered time-linear coefficients 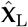 and the centered time-periodic coefficients 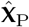. The time-linear and time-periodic PCs are themselves defined by the columns of the 4 × 4 coefficient matrix **V**_L_ and the 36 × 36 coefficient matrix **V**_P_. Because the coefficient matrices are centered in this analysis, it follows that the PCs describe each wingbeat oscillation as a deviation from the mean wingbeat oscillation, with the first PC describing the strongest coupling of the kinematic variables over all of the wingbeats, the second PC the next strongest coupling, and so on. The data and code required to perform this decomposition are provided in MATLAB format as Supplementary Data 1, and the resulting matrices are also provided separately as text files in Supplementary Data 2.

There are two key reasons for applying FPCA here. The first is to gain insight into how the wing kinematics vary between wingbeats, noting that the eigenvectors contained in the leading columns of the PC coefficient matrices **V**_L_ and **V**_P_ describe the dominant couplings of the time-periodic and time-linear terms, respectively. The second is to reduce the complexity of the functional model of the wing kinematics through model reduction, noting that the magnitude of the eigenvalues contained in the matrices ***Λ***_L_ and ***Λ***_P_ describes the amount of variation explained by the corresponding PC eigenvector. In fact, as we show below, the data matrix **X**_P_ (or similarly the data matrix **X**_L_) can be reconstructed with negligible loss of information as:

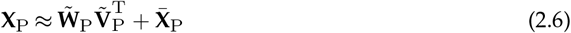

where the tilde notation denotes that the corresponding matrix has been truncated by taking only its first few columns, and where the overbar notation denotes the mean matrix that is removed by centering the data, where 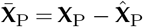. The data and the code required to perform this reconstruction are provided in MATLAB format as Supplementary Data 1

## 3. Results

### (a) FPCA provides a high degree of data reduction

The primary result of the FPCA is a compact representation of the entire dataset, in which the kinematics of each wingbeat are represented as a perturbation from the kinematics of the mean wingbeat. These perturbations are described by the PC scores contained in the 2*N*-by-40 matrix **W** = [**W**_L_ | **W**_P_] where each of the 2*N* rows provides information on one of the *N* = 26, 541 wingbeat pairs measured for either the left or the right wing. The physical meaning of the PC scores is given by the corresponding PC coefficient matrices **V**_L_ and **V**_P_, of dimension 4 × 4 and 36 × 36, respectively. Given knowledge of the time history of the mean wingbeat, it follows that the complete time history of every wingbeat can be reconstructed at full fidelity from its score on each of the 40 time-linear and time-periodic PCs in conjunction with the information contained in the PC coefficient matrices.

In practice, the dimension of the data can be further reduced by considering only the most informative subset of the PCs. For example, the first 20 time-periodic PCs are sufficient to explain > 99% of the measured variation about the mean wingbeat in the time-periodic components of all *N* = 26, 541 wingbeat pairs (Figure 2). There is therefore minimal loss of information on the time history of the wingbeat associated with truncating the 2*N*-by-36 matrix **W**_P_ to a 2*N*-by-20 matrix 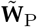 (Figure 3). It is important to note that the remaining < 1% of the variation in the measured kinematics is likely to represent noise rather than real underlying variation in the wing kinematics. By the same token, only the first 2 time-linear PCs are needed to explain > 99% of the measured variation in the linear transition components of all *N* = 26, 541 wingbeats (Figure 2). In this case, however, it is appropriate to use all of the columns of the 2*N*-by-4 linear transition matrix **W**_L_ to avoid the discontinuities in the kinematics that would otherwise be present at the transitions between successive wingbeats.

**Figure 2.**
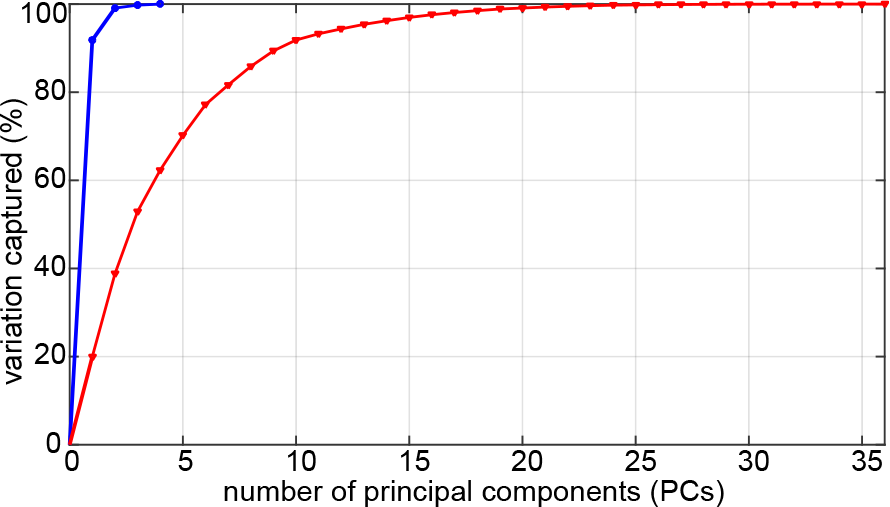
Explanatory power of the principal components (PCs). Graph plots cumulative percentage of between-wingbeat variation in the time-linear and time-periodic wing kinematics that is explained by the time-linear (blue) and time-periodic (red) PCs over all *N* = 26, 541 wingbeat pairs for both the left and right wings.

**Figure 3.**
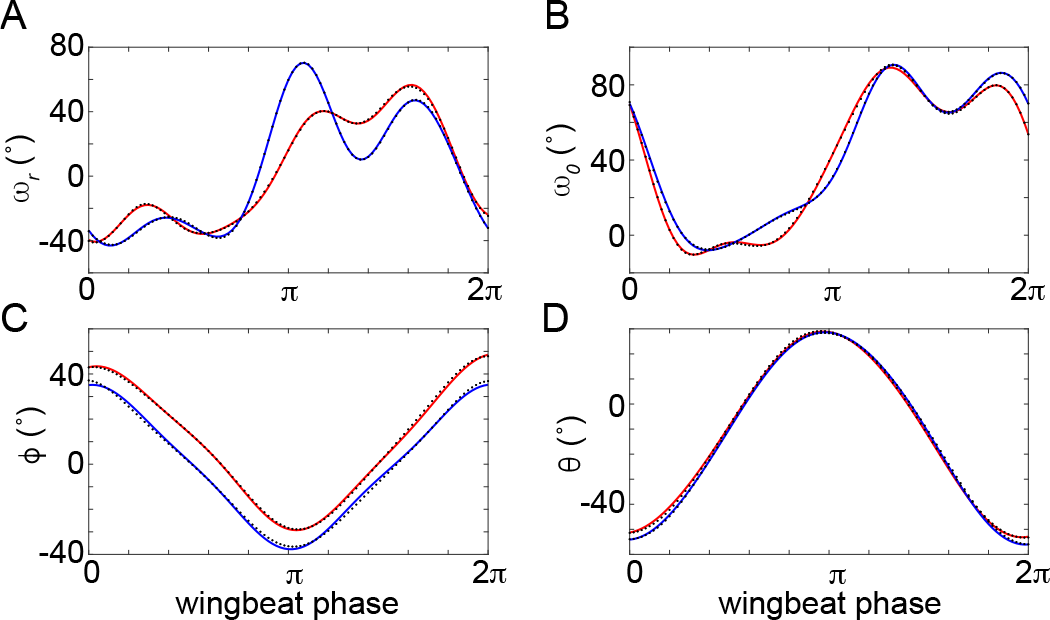
Kinematic reconstruction of a representative wingbeat pair. Time history of the four primary kinematic variables for the left (red) and right (blue) wings, shown for a single wingbeat chosen at random from the first flight sequence, and plotting (A) twist gradient (*ω*_*r*_), (B) twist constant (*ω*_0_), (C) stroke angle (*ϕ*), and (D) deviation angle (*θ*). Dots denote the full fidelity kinematics obtained by evaluating the splines fitted to the raw measurements at 101 evenly spaced points through the wingbeat on the interval *φ* ∈ [0, 2*π*]. Solid lines show the reduced-order reconstructions obtained by using only the first 20 time-periodic PCs and all 4 time-linear PCs. This is sufficient to reconstruct > 99% of the measured variation in the wingbeat kinematics over all *N* = 26, 541 wingbeat pairs. Moreover, because this variation is expressed with respect to the mean wingbeat kinematics, which provide an exact time-varying baseline for the reconstruction, the actual time history of any individual wingbeat will be captured at much better than 99% accuracy. See Supplementary Video 2 for an animated version of this figure showing the effect of varying the number of time-periodic PCs used in the reconstruction.

Putting all this together, it follows that almost all of the time-continuous variation in wing kinematics over all 2*N* = 53, 082 wingbeat pairs can be captured by the 2*N*-by-20 matrix 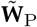 and by the 2*N*-by-4 matrix **W**_L_. The extent of the approximation involved is illustrated for one randomly reconstructed wingbeat in Figures 3 and 1 (see also Supplementary Videos 1–2). Further reduction of the dimensionality of the problem is of course possible, depending on the intended use and required accuracy of the reduced order modelling (see Discussion). This low-dimensional representation of the data is possible for only two reasons: first, because the wingbeat kinematics are quasi-periodic, with only a few significant harmonics; and second, because the dominant PCs capture the natural couplings inherent to the insect’s wing kinematics. These natural couplings manifest themselves over the entire dataset as statistical correlations between the orthogonal basis functions that were used to describe each wingbeat (see Eq. 2.1), and they are fully described by the columns of the PC coefficient matrices 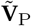 and **V**_L_. Thus, not only does the identification of the most informative set of PCs provide a high degree of data reduction; it also provides insight into the natural couplings that arise from the actuation, dynamics, and control of the insect’s wings.

### (b) FPCA identifies repeatable couplings in the wing kinematics

To check the reliability of the couplings identified by the FPCA, we ran a validation analysis on centered subsets of the data matrix **X**_P_, comparing the PC coefficients for these subsets with one another or with those already obtained. We first checked the similarity of the PC loadings identified independently for the left and right wings, reasoning that these ought in principle to be self-similar. We defined the mean absolute correlation coefficient matrix **R** for this case as the matrix with entries *R*_(*i,j*)_ corresponding to the absolute correlation coefficient between the *i*^*th*^ and *j*^*th*^ columns of the time-periodic PC coefficient matrices of the left and right wings. As expected, this matrix was nearly diagonal (Figure 4A), indicating that the identified couplings do indeed have a high degree of self-similarity for the left and right wings. We next ran a resampling analysis in which we repeated the FPCA 100,000 times on random subsamples each containing 10% of the 2*N* = 53, 082 rows of the data matrix **X**_P_, drawn without replacement. We defined the mean absolute correlation coefficient matrix **R** for this case as the matrix with entries *R*(*i,j*) corresponding to the mean of the absolute correlation coefficients between the *i*^*th*^ column of the full time-periodic PC coefficient matrix and the *j*^*th*^ column of this matrix for each of the random subsamples. This matrix was again nearly diagonal (Figure 4B), indicating that the identified couplings have a high degree of consistency across different subsamples of the dataset. It follows that the identified couplings describe repeatable features of the data, and that it would also have been possible to identify these using a sample an order of magnitude smaller than the *N* = 26, 541 wingbeat pairs analysed here.

**Figure 4.**
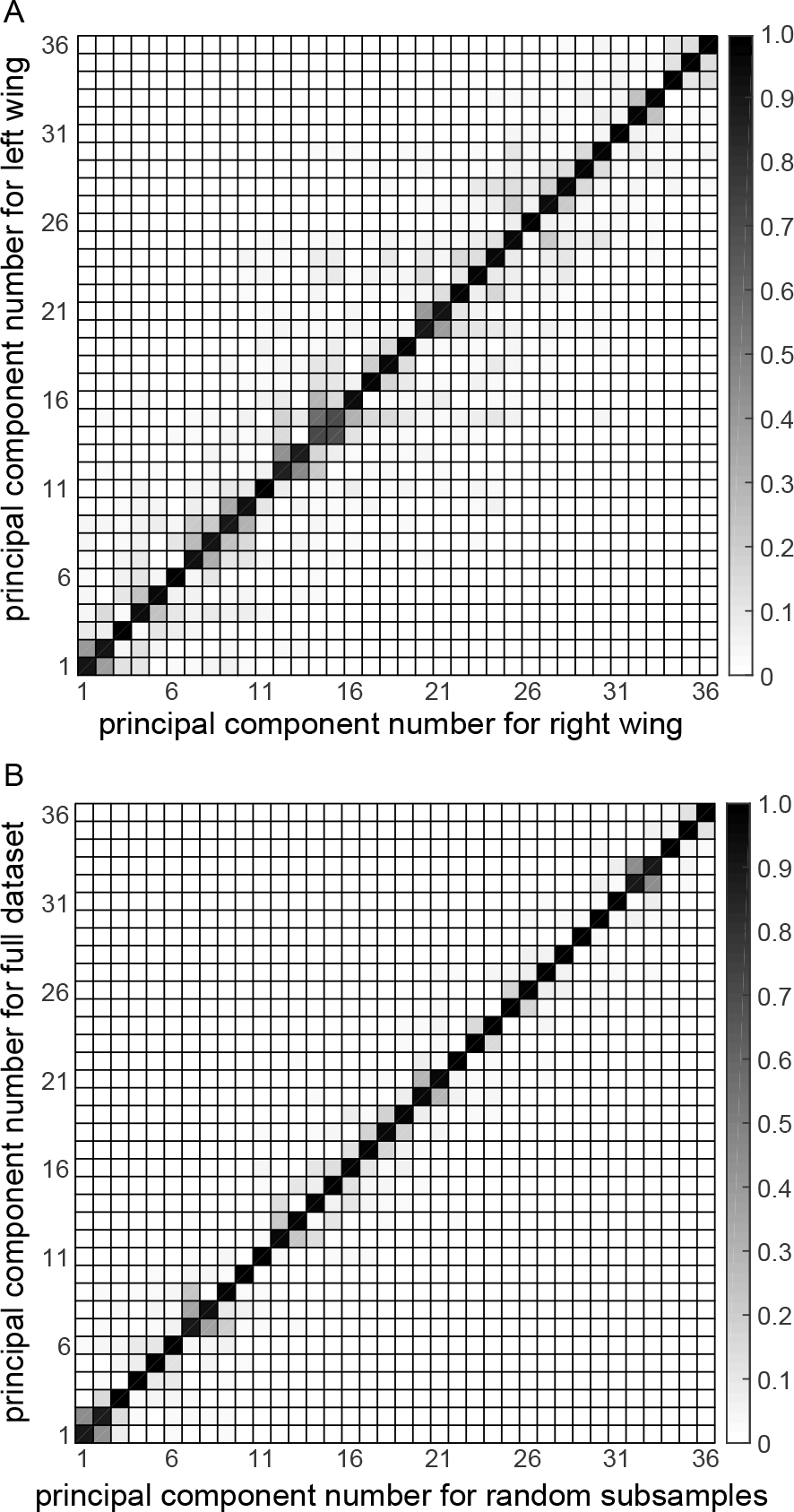
Repeatability of the identified principal component (PC) couplings. Absolute correlation coefficient matrix for the 36 time-periodic PCs, ordered according to the proportion of the total variation that each PC explains. The grayscale level of the cells corresponds linearly to the absolute value of the correlation coefficient, calculated as: (A) the absolute correlation coefficient between the PC coefficient vectors for the left and right wings; and (B) the mean of the absolute correlation coefficient between the PC coefficient vectors identified for the full dataset, and for each of 100,000 random subsamples comprising 10% of the full dataset. Note that each correlation coefficient matrix is diagonally dominant, indicating that similar sets of PCs are identified for the left and right wings, and for random subsamples of the data as compared to the full dataset. The couplings that the PC coefficient vectors describe are therefore repeatable, hence physically meaningful, features of the data.

### (c) Wing twist variation dominates the kinematic couplings

We next sought to characterize the variation in wing kinematics described by each PC. All of the most important time-periodic and time-linear PCs are dominated by variation in the wing twist variables, *ω*_0_ and *ω*_*r*_, coupled with a much smaller degree of variation in the stroke angle *ϕ*, and an even smaller degree of variation in the deviation angle *θ* (Figure 5). This partly reflects the fact that the wingbeat phase *φ* was always determined with reference to the wing tip kinematics (see Materials and methods), which means that the stroke angle and deviation angle have less scope to vary between wingbeats at any given phase than do the wing twist variables. Even so, the fact that all of the most important PCs are dominated by variation in wing twist indicates that the phasing of this wing twist is quite variable with respect to the phase of the wing tip kinematics (see Discussion).

**Figure 5.**
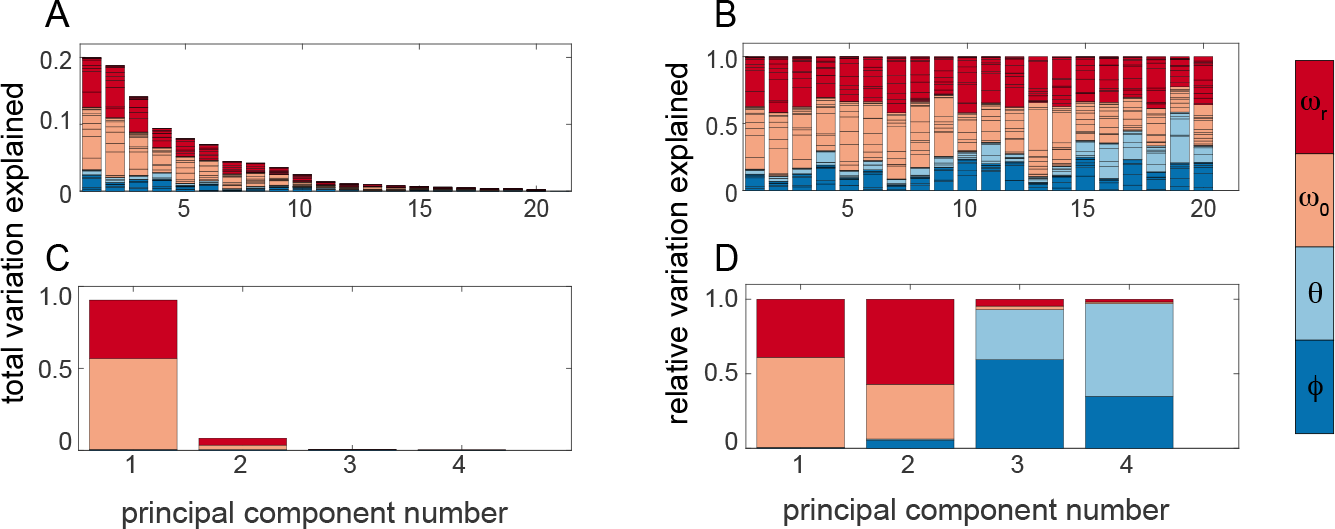
Contribution of the four primary kinematic variables to the dominant principal component (PC) couplings. (A,B) Contributions to time-periodic PCs 1–20; higher-numbered PCs not shown, on the basis that individually they have trivial explanatory power. (C,D) contributions to time-linear PCs 1–4. Each vertically stacked bar represents a single PC coefficient vector; each subdivision shows the magnitude of a PC coefficient corresponding to one of the terms in the original Fourier series representation (see Eq. 2.1). Subdivisions are coloured according to the primary kinematic variable to which they refer: *ϕ* (dark blue), *θ* (light blue), *ω*_0_ (light red), or *ω*_*r*_ (red). In (A,C), the PC coefficients are normalized such that the total height of the bar for each PC is equal to the proportion of the total variation that it explains. In (B,D), the coefficient are normalized so that all of the bars have the same total height. The former gives an indication of how much of the overall variation in wing kinematics is attributable to each primary kinematic variable; the latter allows direct comparison of the relative importance of the different primary kinematic variables to each PC. Note the especially strong contribution of the wing twist variables to all of the dominant PCs (i.e. lowest numbered PCs).

### (d) Time-domain analysis of the dominant kinematic couplings

We visualised the dominant kinematic couplings by perturbing the time-history of the four primary kinematic variables with respect to the mean wingbeat, using the PC coefficients contained in the leading columns of the matrices 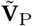 and **V**_L_ (Figures 6 and 7). In doing so, we used the standard deviations (SDs) of the PC scores across all 2*N* = 53, 082 left and right wingbeats (i.e. the column-wise SDs of 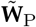 and **W**_L_) to weight the PC coefficients, thereby revealing the typical range of variation in the four time-varying primary kinematic variables that is attributable to each PC (Figure 6). This analysis shows, for example, that time-periodic PC1 involves continuous changes in wing twist that are most pronounced through the upstroke, coupled with large changes in stroke angle that are greatest at stroke reversal (Figure 6a). In contrast, the variation in wing twist that characterises time-periodic PC2 throughout the stroke is less strongly coupled with variation in the wing tip kinematics (Figure 6b). Time-periodic PC3 involves pronounced variation in stroke angle at dorsal stroke reversal, coupled with changes in wing twist throughout the wingbeat (Figure 6c).

**Figure 6.**
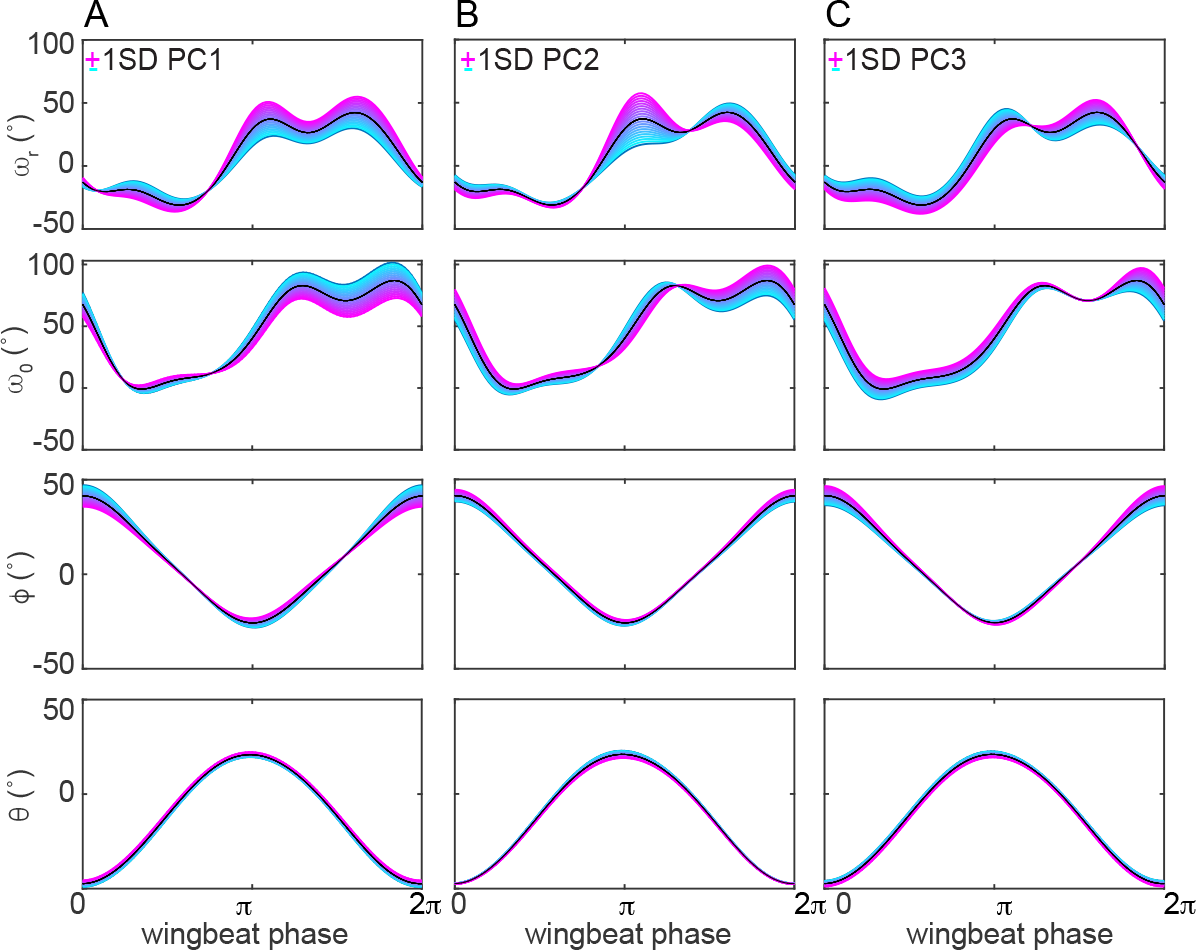
Time-domain representation of the dominant time-periodic principal components (PCs). The plots show how the time histories of the four primary kinematic variables are affected by perturbing the kinematics of the mean wingbeat ±1SD in each of the first 3 time-periodic PCs, where SD denotes the standard deviation of the scores for that PC over all wingbeats: (A) PC1; (B) PC2; (C) PC3. Black lines plot the time history of the stroke angle (*ϕ*), deviation angle (*θ*), twist gradient (*ω*_*r*_), and twist constant (*ω*_0_); cyan and magenta regions of the graph display the effect of a ±1SD perturbation.

**Figure 7.**
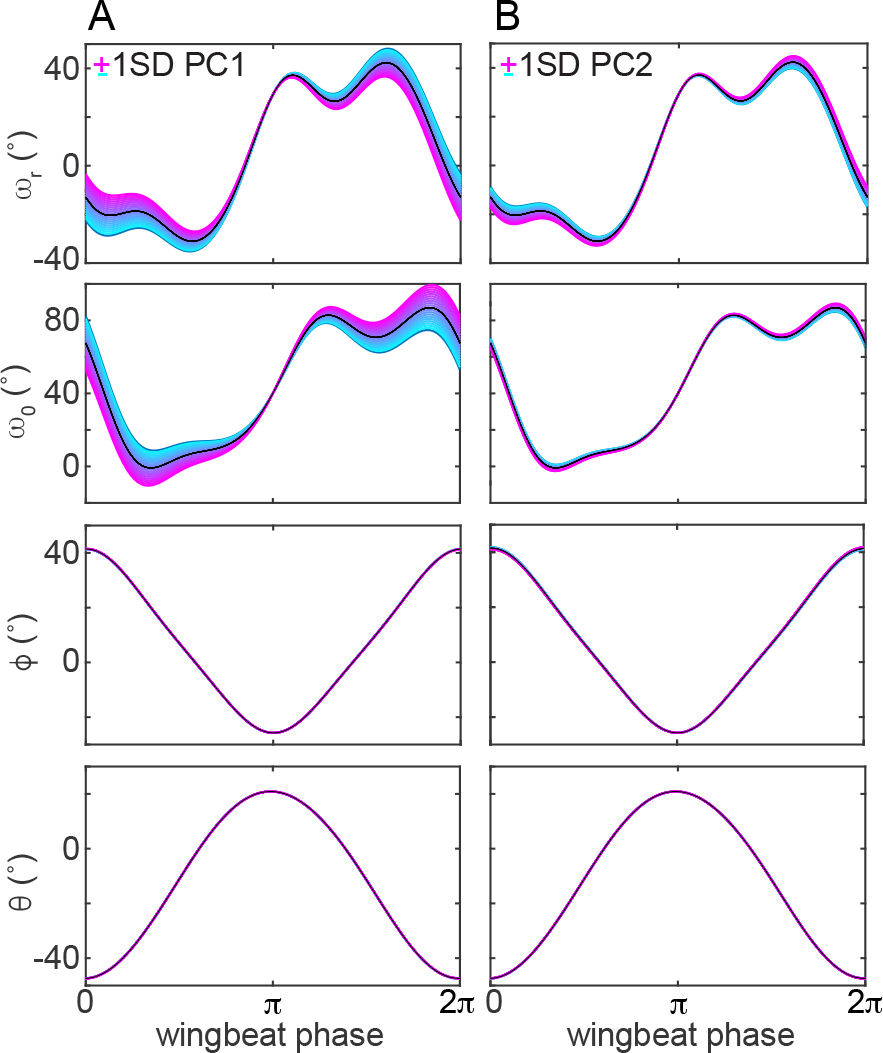
Time-domain representation of the dominant time-linear principal components (PCs). The plots show how the time histories of the four primary kinematic variables are affected by perturbing the kinematics of the mean wingbeat ±1SD in each of the first 2 time-periodic PCs, where SD denotes the standard deviation of the scores for that PC over all wingbeats: (A) PC1; (B) PC2. Black lines plot the time history of the stroke angle (*ϕ*), deviation angle (*θ*), twist gradient (*ω*_*r*_), and twist constant (*ω*_0_); cyan and magenta regions of the graph display the effect of a ±1SD perturbation.

The corresponding analysis of the time-linear PCs shows that time-linear PC1 involves opposing changes in twist gradient and twist constant (Figure 7a), whereas time-linear PC2 involves the same direction of change in both wing twist parameters (Figure 7a). In both cases, the associated changes in wing tip kinematics are negligible (Figure 7), which makes the inclusion of time-linear PCs 3–4 particularly important to avoiding discontinuities in the wing tip kinematics at the transitions between wingbeats. Taken together, the first three time-periodic PCs are sufficient to capture more than half the variation in the time-periodic wing kinematics between wingbeats, whilst the first two time-linear PCs already capture almost all of the variation in the time-linear wing kinematics between wingbeats (Figure 2). It follows that Figures 6–7 contain information on the kinematic couplings that together explain the majority of the variation between wingbeats. A similar exercise can be carried out for as many PCs as may be required.

### (e) Phase-domain analysis of the dominant kinematic couplings

It is already evident from the time domain plots in Figure 6 that the time-periodic PCs each manifest themselves most strongly at characteristic phases of the stroke cycle. The detailed phase relationships of these time-periodic kinematic couplings are even more clearly visible in the corresponding phase portraits. To understand the effects of these perturbations, it is helpful first to consider the mean wingbeat kinematics, which form the baseline for the perturbed wingbeat kinematics (Figure 8). The mean wingtip trajectory is similar on both the upstroke and the downstroke, as can be seen by inspection of the phase portrait of stroke angle against deviation angle (Figure 8A). In contrast, the wing twist variables follow an opposed path on the upstroke as compared to the downstroke (Figure 8B), which reflects the fact that the wing is inverted on the upstroke so that its aerodynamic angle of attack is similar on both half strokes. The complex, looped nature of this path reflects the harmonic content that is present in the wing twist through the mean wingbeat (cf. Figure 6).

**Figure 8.**
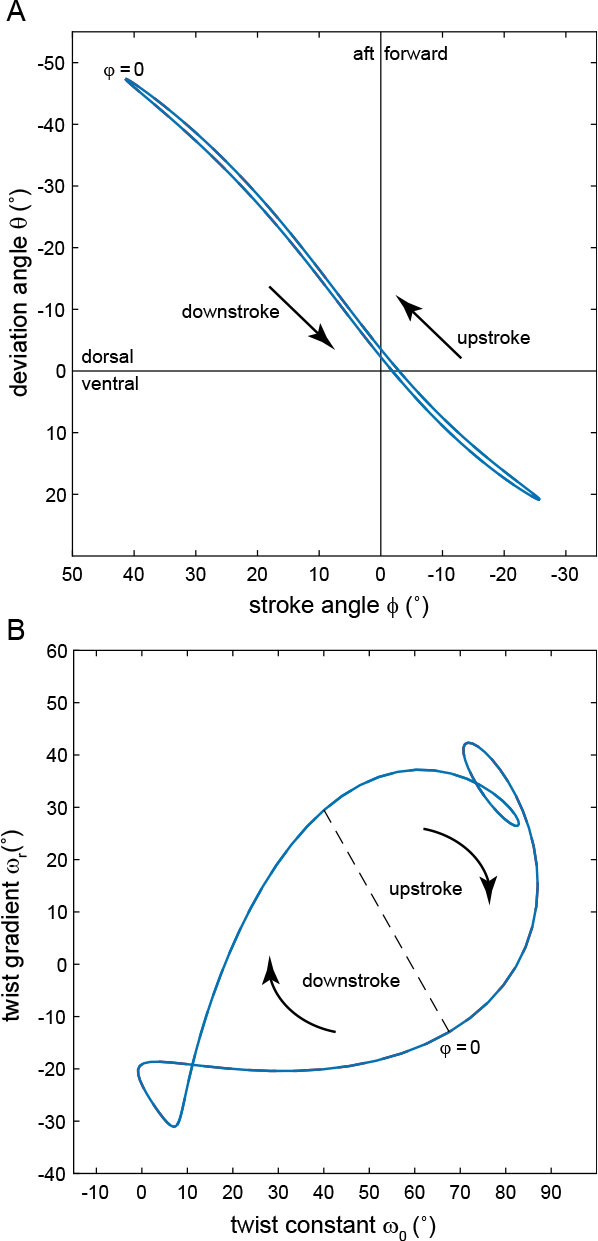
Phase portraits of the mean wingbeat kinematics. (A) Wing tip kinematics, plotting deviation angle (*θ*) versus stroke angle (*ϕ*); and (B) wing twist kinematics, plotting twist gradient (*ω*_*r*_) versus twist constant (*ω*_0_). Zero phase (*φ* = 0) denotes the start of the downstroke. Note that as *ϕ* and *θ* are the spherical coordinates of the wing tip, the phase portrait in (A) may be interpreted as an equirectangular projection of the wingtip trajectory; there is no straightforward physical interpretation of the phase portrait in (B), but both phase portraits show clearly how the primary kinematic variables are coupled in the mean wingbeat.

The detailed effect of each time-periodic PC is described by the change it produces in the shape, position, or orientation of the phase portrait. The kinematic effect of any given PC is reversible, so we adopt the convention of using parentheses to show how the different elements of the PC coefficient vector are coupled. For example, under time-periodic PC1, a steepening (shallowing) of the stroke plane is associated with a shortening (lengthening) of the stroke path (Figure 9A), but the same steepening (shallowing) of the stroke plane could in principle have been associated with a lengthening (shortening) of the stroke path under an alternate kinematic coupling. This steepening (shallowing) and shortening (lengthening) of the stroke path under time-periodic PC1 results in a marked decrease (increase) in the overall amplitude of the wingbeat, and is accompanied by compression (expansion) of the path traced by the wingtip on the lower half of the stroke (Figure 9A). These changes in wing tip kinematics are associated with an increase (decrease) in the amplitude of the oscillatory variation in wing twist gradient, and with a decrease (increase) in the amplitude of the oscillatory variation in wing twist constant (Figure 10A). The corresponding changes in wing pitch are most pronounced at the points of dorsal and ventral stroke reversal, when the distance between the equivalent points on the reference and perturbed limit cycles is greatest (Figure 10A). Overall, the effect of perturbing PC1 is to steepen (shallow) the stroke plane whilst decreasing the stroke amplitude, and increasing (decreasing) the degree to which the wing’s local pitch angle varies as a result of the coupled changes in wing twist constant and wing twist gradient.

**Figure 9.**
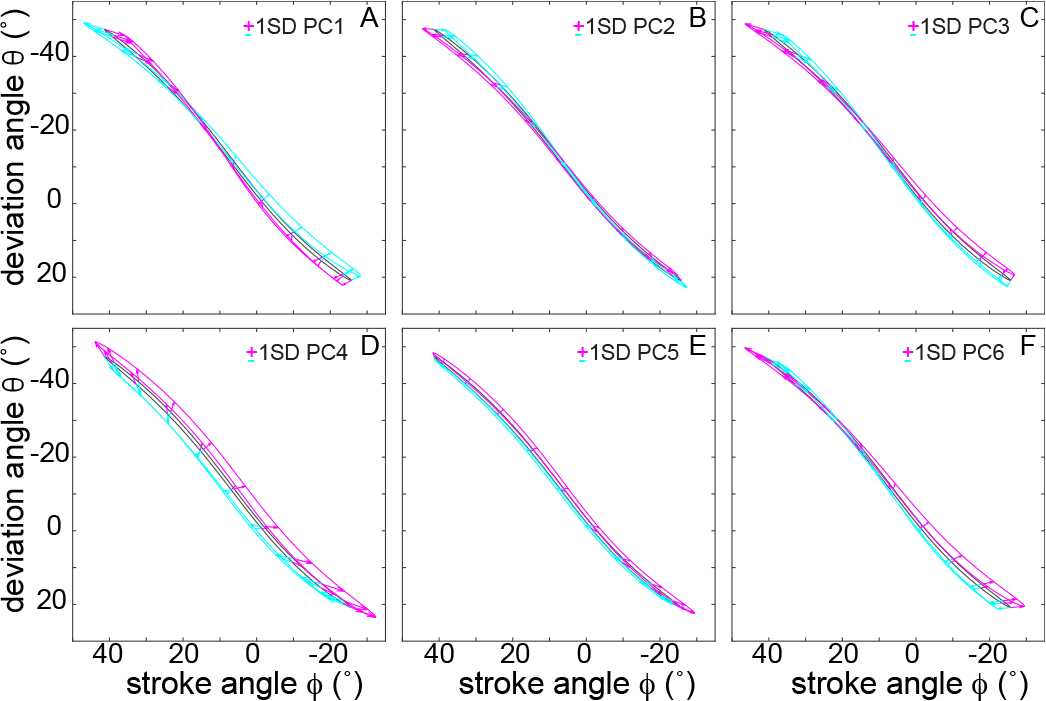
Phase-portraits showing the effects of the dominant time-periodic principal components (PCs) on the wing tip kinematics. Black line plots the phase portrait of the mean wingbeat (see Figure 8A) for deviation angle (*θ*) versus stroke angle (*ϕ*); cyan and magenta lines display the effect of a ±1SD perturbation in each PC, with arrows connecting points of equivalent phase. Note that perturbing some PCs (e.g. PC1) results in a change of the slope of the wingtip trajectory, whereas perturbing others results in a wholesale displacement of the wingtip trajectory (e.g. PC5) or a change in the width of the loop swept (e.g. PC4).

**Figure 10.**
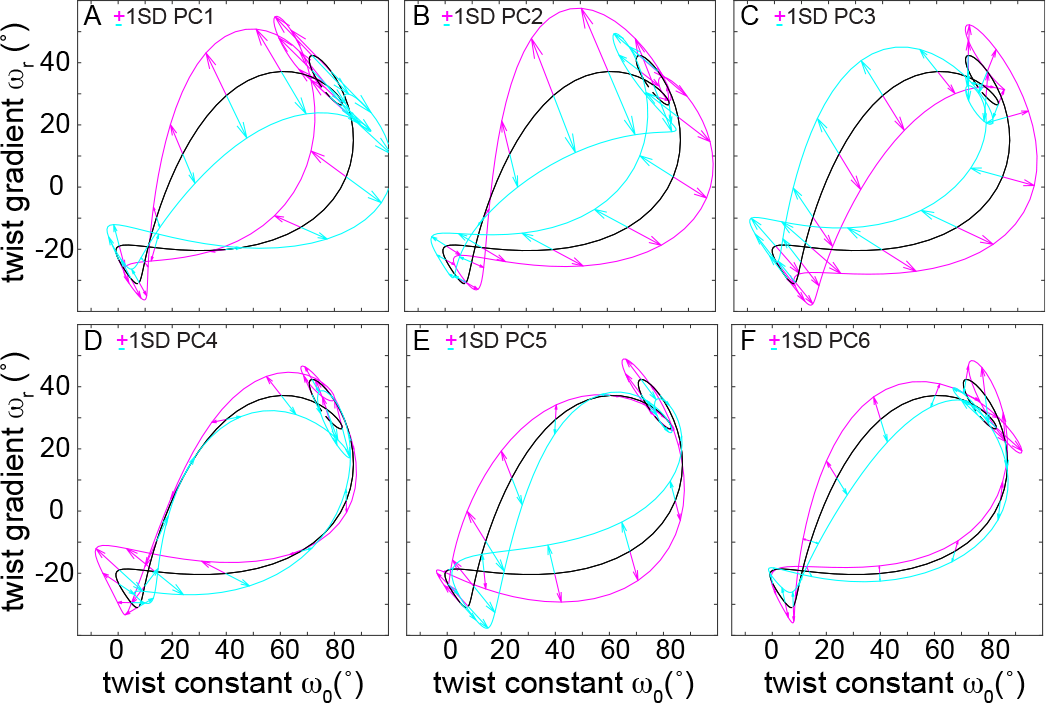
Phase-portraits showing the effects of the dominant time-periodic principal components (PCs) on the wing twist kinematics. Black line plots the phase portrait of the mean wingbeat (see Figure 8B) for twist gradient (*ω*_*r*_) versus twist constant (*ω*_0_); cyan and magenta lines display the effect of a ±1SD perturbation in each PC, with arrows connecting points of equivalent phase. Note that the different PCs affect different phases of the wingbeat to a greater or lesser degree.

In contrast, time-periodic PC2 describes shallowing (steepening) of the stroke plane, which is most pronounced through the upper half of the stroke, and which is accompanied by only minimal changes in the shape of the wingtip path (Figure 9B). This shallowing (steepening) of the stroke plane is accompanied by a coupled increase (decrease) in the amplitude of the oscillatory variation in both the wing twist constant and wing twist gradient (Figure 10B). Again, the resulting changes in wing pitch angle are most pronounced at the points of dorsal and ventral stroke reversal. In summary, the general effect of perturbing PC2 is to shallow (steepen) the stroke plane, whilst greatly increasing (decreasing) the extent of the variation in the wing’s pitch angle through the wingbeat through its effects on both the rigid-body rotation and torsional deformation of the wing. In contrast, time-periodic PC3 describes changes in wingtip trajectory (Figure 9C) that look geometrically to similar those of PC1 (Figure 9A), albeit that its PC coefficient vector happens to be oppositely signed. However, whereas the changes in the wingtip trajectory under PC1 are associated with opposed changes in the amplitude of the oscillatory variation in wing twist constant and gradient, in PC3 they are associated with a wholesale shift in the position of the orbit describing the variation in wing twist constant and gradient through the wingbeat (Figure 10C). In summary, the general effect of perturbing PC3 is to cause a shallowing (steepening) of the stroke plane, whilst decreasing (increasing) the extent of the wing’s torsional deformation and increasing (decreasing) its pitch angle at the base.

Time-periodic PCs 1-3 are already sufficient to capture > 50% of the variation in the time-periodic wing kinematics between wingbeats, whilst time-periodic PCs 4-6 collectively explain another 30% of this variation. Time-periodic PCs 4-6 are each associated with lesser changes in wing twist constant and wing twist gradient than PCs 1-3 (Figures 10D-F; cf. Figures 10A-C), but the associated changes in wingtip trajectory are at least as large (Figures 9D-F; cf. Figures 9A-C). Focussing on these changes in wingtip trajectory, time-periodic PCs 4 and 5 are both associated with a prominent increase (decrease) in stroke angle throughout the lower half of the stroke, leading to a forward (aft) displacement of the position of the wingtip at ventral stroke reversal (Figures 9D-E). They are also both associated with an increase (decrease) in deviation angle at the upper part of the stroke, leading to a dorsal (ventral) displacement of the wingtip at dorsal stroke reversal, which is much more prominent for PC4 (Figure 9D). PC6, in contrast, involves a steepening (shallowing) of the stroke plane and an expansion (compression) of the path traced by the wingtip on the lower half of the stroke (Figure 9F), which resembles closely the pattern observed for PC3 (Figure 9C).

## 4. Discussion

The time-periodic and time-linear PCs that we have identified provide a compact decomposition of the natural couplings that are inherent in the wing kinematics of manoeuvring hoverflies. Thus, any steady-state oscillation of the wing can be characterised as a perturbation of the reference wingbeat along the directions defined by the time-periodic PCs that we have identified. Any non-steady oscillation, in which the wing ends its stroke in a different kinematic state to that in which it began, can be described by superimposing the time-linear PCs that we have identified. The approach that we have developed here can therefore be considered a model-based approach to data reduction that reflects the characterisation of insect flight as involving a form of limit cycle control [4,6,7].

The observation that only around half the PCs are needed to capture > 99% of the measured variation in the wing kinematics relates partly to the fact that the higher harmonics of some of the fitted Fourier series are unimportant, and are effectively dropped by the FPCA. For example, to explain > 99% of the between-wingbeat variation in each of the four kinematic variables it is only necessary to include up to the 1st harmonic for stroke angle, up to the 3rd harmonic for deviation angle, and up to the 4th harmonic for twist constant. All 5 fitted harmonics are needed to explain > 99% of the between-wingbeat variation in twist gradient. Even so, reducing the order of the Fourier series independently in this way would still leave us with a total of 34 fitted coefficients, compared to the 24 PCs used to reconstruct the data in Figure 3. It follows that a considerable part of the data reduction that we have achieved through the FPCA must be attributable to the couplings between the kinematic variables that are described in the Results above. In any case, it is clear that the overall result of the FPCA is an exceedingly compact representation of the wing kinematics.

In this compact representation, all of the time-continuous kinematic variation over our entire dataset of *N* = 26, 541 wingbeat pairs is summarised for both the left and the right wings by the 2*N*-by-40 matrix **W** at effectively full fidelity, and by the 2*N*-by-24 matrix 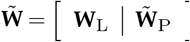 at > 99% fidelity. Lower-dimensional representations are possible using fewer PCs, depending on how high a degree of approximation is acceptable in the data. The only pieces of information that are required in addition to the matrix **W** (or 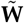) are: (i) the corresponding 4-by-4 matrix **V**_L_ and the 36-by-36 matrix **V**_P_ (or its truncated submatrix 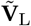); (ii) the vector of length 40 containing the Fourier series coefficients for the mean reference wingbeat; (iii) the vector of length 2*N* containing the period of each wingbeat; and (iv) the details of the wing’s planform. All these data are published here as Supplementary Data 1, to enable this dataset to serve as the basis for future analyses using computational fluid dynamics (CFD) or other aerodynamic modelling approaches. Because the entire dataset is uniquely defined in functional form, it can be used reliably for the cross-validation of different flow solvers and meshing approaches, because different researchers will be able to generate the same, fully-repeatable wing kinematics with whatever meshing approach they use.

The natural couplings that the PCs identify relate fundamentally to the control that an insect can exert over its wing kinematics. Specifically, whilst the insect may not be able to control every PC independently of every other, and whilst the PCs necessarily capture inter- as well as intra-individual variation in wing kinematics, the subset of dominant PCs must collectively span the entire range space of the insect’s flight control system—insofar as this is represented in the data. In other words, the dominant PCs are sufficient to contain all of the wing kinematics that the insect actually uses. Conversely, the subset of PCs that prove unimportant in our analysis must describe all of the geometrically feasible kinematic couplings that for whatever reason the insect cannot—or does not—use. The overall conclusion that the PCs tell us something fundamental about *Eristalis* flight control is confirmed by the fact that the FPCA extracts repeatable features from the data. We have demonstrated this in two distinct ways. First, we have shown that essentially the same set of PCs is identified for each insect if the analysis is run on the left and right wings separately. Second, we have shown that the same set of principal components is repeatably identified by running the analysis on 100,000 random subsamples of the data. It follows that by identifying the dominant couplings in the wing kinematics, the PC decomposition provides a natural basis in which to describe how an insect controls its flight.

All of the dominant time-periodic PCs can be interpreted in terms of their effects on the familiar summary kinematic variables including stroke amplitude, stroke plane angle, and the timing of wing rotation. However, the full descriptions of the most important PCs that are provided in Figures 6–7 and 9–10, and which are implicit in the PC coefficient vectors themselves (Supplementary Data 1), show that the reality is much richer than can be captured by these summary variables alone. For example, a steeper stroke plane angle is typically associated with a lower stroke amplitude (Figure 9A,C,F), but is also associated with changes in the shape of the path traced by the wingtip, and with coupled changes in the detailed time history of the wing’s twist. Indeed, the overriding conclusion that emerges from this analysis is that the most variable components of the wing kinematics are those involving wing twist, and comprise complex, high-order changes in the phasing of the wing’s twisting motion throughout the stroke (Figure 10). Whilst this variability may partly reflect the difficulty of regulating the passive torsional modes of the wing through only the application of a pitching torque at the wing root, the fact that specific patterns of variation in wing twist are repeatably coupled to specific patterns of variation in wing tip trajectory clearly indicates that a large portion of this variability must indeed be under the insect’s direct control. Moreover, the nature of these couplings is such that the dominant PCs all involve changes in the phasing of wing pronation and supination relative to dorsal and ventral stroke reversal, respectively (Figure 10). This is interesting, because it is well known that the aerodynamic forces are highly sensitive to the timing of wing pronation and supination relative to the timing of dorsal and ventral stroke reversal [25,26]. This already suggests that the PCs which capture the most variation in the wing kinematics are likely to be important aerodynamically, as does the broader observation that the fine details of the wing’s motion can have large effects on aerodynamic force production [18]. A detailed analysis of the aerodynamic effects of the PCs will be provided elsewhere.

In conclusion, there are two complementary ways of interpreting the physical meaning of the PC decomposition. If the aim is to understand how muscle actions control wing kinematics, then the dominant PCs define an appropriate basis for observing the kinematic output of the insect’s neuromuscular control system. Alternatively, if the aim is to understand how wing kinematics control flight dynamics, then the PCs define an appropriate basis for observing the kinematic inputs to the insect’s flight dynamics. Hence, the PCs that we have identified may be used to define either a natural state vector for the muscle actuation system, or a natural control input vector for the flight control system. This is in contrast to the more conventional approach in insect flight dynamics of taking the kinematic variables that a human observer might otherwise define—stroke amplitude, stroke plane angle, and so on—as motor outputs or control inputs. Future work will make use of the basis provided by the reduced set of PCs to build physically and biologically meaningful state-space models for systems-level analyses of observability and controllability.

## Data Accessibility

The supplementary data and code supporting this article are available through figshare: https://doi.org/10.6084/m9.figshare.10269614

## Authors’ Contributions

SW collected and provided the published dataset on which this analysis is based, wrote code, and prepared figures. IN and GT analysed the data, wrote code, prepared figures, and wrote the paper. All authors gave final approval for publication and agree to be held accountable for the work performed therein.

## Acknowledgements

We thank Sean Humbert and Holger Krapp for many useful discussions, and thank Marco Klein Heerenbrink for commenting on a draft of this manuscript.

## Supplementary Data 1

This zipped folder contains all of the data used in the analysis in the main paper, together with code for implementing the various steps of the analysis, both in MATLAB format. Running the m-files in sequence produces a reduced-order approximation and reconstruction of the data. See README file and comments in the various m-files for further details.

## Supplementary Data 2

This zipped folder contains plain text files corresponding to the data matrices **X**_L_ and **X**_P_ in units of degrees, together with the results of the functional principal components analysis on these data matrices, comprising: (i) the mean matrices 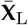 and 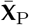; (ii) the PC score matrices **W**_L_ and **W**_P_; (iii) the PC coefficient matrices **V**_L_ and **V**_P_; and (iv) the vectors of eigenvalues on the diagonals of the matrices ***Λ***_L_ and ***Λ***_P_. The same folder also contains a vector giving the period of each wingbeat (ms), a vector giving the wing length for each wingbeat (m), and matrices containing the chordwise and spanwise coordinates of 100 points on both the leading and trailing edges of a representative wing outline of *Eristalis*. These coordinates are expressed in dimensionless form, and may be put in dimensional form by multiplying them by the wing length. The data matrices and PC score matrices of length 2*N* = 53, 082 are partitioned such that the first *N* = 26, 541 rows correspond to the left wingbeats, and the next *N* = 26, 541 rows correspond to the right wingbeats. See Materials and methods for details and nomenclature.

## Supplementary Video 1

Kinematic reconstruction of a randomly selected wingbeat, showing the effect of increasing the number of time-periodic principal components (PCs) used in the reconstruction from 1 to 36. Each reconstruction uses all of the time-linear PCs to ensure continuity at the transitions between wingbeats. Left column shows wing coloured by its local pitch angle; right column shows wing coloured by the difference between its local pitch angle and that of the full-fidelity reconstruction from the Fourier series analysis.

## Supplementary Video 2

Animation of Figure 3 showing the effect of varying the number of time-periodic principal components (PCs) used in the reconstruction from 1 to 36. Each reconstruction uses all of the time-linear PCs.

